# Vestibular Perturbation Disrupts Source and Reality Monitoring

**DOI:** 10.64898/2026.08.07.743488

**Authors:** C. M Bates, L Lingawi, E Perry, M Gallagher, A. K Martin

**Affiliations:** School of Psychology, The University of Kent, UK; Kent Medway Medical School, UK

**Keywords:** noisy galvanic vestibular stimulation (nGVS), reality monitoring, source monitoring, depersonalisation/derealisation, self–other processing

## Abstract

The vestibular system has been implicated in several aspects of self-representation, including bodily self-location, yet its role in episodic source and reality monitoring remains unclear. Moreover, symptoms of depersonalisation and derealisation (DPDR) have been associated with both vestibular dysfunction and altered source and reality monitoring. We tested whether noisy galvanic vestibular stimulation (nGVS) modulates source and reality monitoring during a word-encoding paradigm and whether this increased symptoms of DPDR. Sixty young adults (N = 30 active nGVS, N = 30 sham) encoded words that varied by context (spoken vs. imagined), agentic source (self vs. other), and valence (positive vs. threat) before completing source/reality judgments. Active nGVS reduced overall accuracy relative to sham on both reality and source monitoring. Across conditions, self-referential items were remembered with greater sensitivity compared to other-referential items, and spoken items were recalled with greater sensitivity than imagined items. We additionally show an interaction effect whereby active nGVS reduced accuracy specifically for imagined and threat-valenced items. Trait DPDR did not moderate the effects of stimulation, although higher trait DPDR was associated with a more liberal reality-monitoring response criterion. Results indicate that transient vestibular perturbation via nGVS can impair source monitoring for internalised and threatening stimuli and impairs cognitive distinctions between internally and externally generated events in general, supporting a role for vestibular input in maintaining self-other and reality boundaries in episodic memory.

## 1.1 Vestibular Perturbation Disrupts Source and Reality Monitoring

*Source monitoring* is the cognitive process by which individuals identify the origins of their memories, such as whether an event was tagged to themselves or someone else (Johnson et al., 1993). Within this, Johnson and Raye (1981) additionally describe *reality monitoring*, which comes under the umbrella of source monitoring but specifically references the ability to successfully distinguish between an internal or external event. For instance, remembering whether something happened in real life or in a dream. Although source and reality monitoring have traditionally been explained in terms of cognitive and metacognitive processes (Kuhlmann & Bayen, 2016; Simons et al., 2017), growing evidence suggests that memory is fundamentally embodied, with bodily signals contributing to how experiences are encoded and later retrieved (Ianì, 2019). Interoceptive, proprioceptive, and vestibular information provide continuous cues about the state and position of the body, supporting the integration of self-related information and the distinction between internally generated and externally experienced events (Tsakiris, 2017). The vestibular system has extensive connections with cortical regions implicated in spatial cognition, self-representation, and episodic memory (Hitier et al., 2014), suggesting it may play a previously underappreciated role in memory source attribution. However, despite increasing interest in embodied accounts of memory, the causal contribution of vestibular input to source and reality monitoring remains largely unexplored. The present study addresses this gap by investigating whether noisy galvanic vestibular stimulation (nGVS), a non-invasive technique that modulates vestibular afferent activity, influences performance on source and reality monitoring tasks. In doing so, this study provides one of the first direct tests of whether transient changes in vestibular signalling can alter the processes underlying memory attribution, offering novel insight into the embodied mechanisms supporting source monitoring.

Source and reality monitoring are commonly assessed using episodic memory paradigms that require individuals to remember not only previously encountered information but also the context in which it was acquired (Addante et al., 2024). Source monitoring tasks typically require participants to identify the origin of remembered information, such as whether it was generated by themselves or another person, whereas reality monitoring tasks require discrimination between internally generated experiences (e.g., imagined events) and externally perceived experiences (Johnson & Raye, 1981). Performance on these tasks depends on the successful encoding and retrieval of qualitative characteristics associated with an event, including perceptual, contextual, emotional, and cognitive information that support accurate memory attribution. One paradigm used to assess these processes was developed by Kwon et al. (2022); across two tasks, participants encoded objects associated with either self- or experimenter-generated actions (source monitoring) or imagined versus perceived movements (reality monitoring). Findings from both tasks were aligned, showing participants to be less accurate at attributing internally generated experiences than externally generated experiences. In other words, participants demonstrated an externalisation bias characterised by poorer performance for self- relative to experimenter-generated actions and for imagined relative to perceived movements. As vestibular signals contribute to distinguishing self-generated from externally generated experiences during perception and action (Cullen, 2012; Angelaki & Cullen, 2008; Deroualle & Lopez, 2014), they may also provide an important embodied cue during memory attribution of verbal stimuli.

Emotional valence is known to influence episodic memory and memory phenomenology. For instance, emotional events are typically remembered with greater vividness and subjective recollective experience compared to neutral events (Phelps & Sharot, 2008). Source and reality monitoring rely on the retrieval of qualitative characteristics associated with remembered events, consequently, emotional content may influence monitoring accuracy. Indeed, individuals are less likely to misattribute imagined negative items as perceived, suggesting that emotional arousal strengthens memory for source-specific details (Kensinger & Schacter, 2006). Moreover, threatening stimuli are particularly relevant to bodily self-processing, engaging systems involved in vigilance and defensive responding (Tseng et al., 2023) raising the possibility that vestibular perturbation may differentially affect monitoring for emotionally salient information, related or tagged to the self.

Although source and reality monitoring have traditionally been conceptualised as higher-order cognitive processes, growing evidence suggests that they may depend on embodied representations of the self. Multisensory integration, the process by which the brain combines sensory inputs, has been shown to play a key role in self-other distinction (Tsakiris, 2017).

However, the role of multisensory processing in source and reality monitoring has received little attention. Vestibular signals contribute to a wide range of cognitive functions beyond balance and spatial orientation, including bodily awareness, perspective taking, and self-consciousness (Smith et al., 2005; Miller & Ngo, 2007; Gurvich et al., 2013). Critically, the vestibular system plays a fundamental role in distinguishing self-generated from externally generated motion, meaning that it could well support self-other distinction across other domains (Cullen, 2012; Angelaki & Cullen, 2008; Deroualle & Lopez, 2014).

Clinical observations further underscore the importance of vestibular processing for self-other distinction. For instance, individuals with vestibular dysfunction have difficulties perceiving self-motion and navigation (Britton & Arshad, 2019) and often report reduced feelings of agency and control over the self (Jáuregui-Renaud et al., 2008; Sang et al., 2006).

Importantly, experimental vestibular stimulation in healthy participants can transiently induce similar alterations in bodily self-experience and perceived control (Lopez et al., 2012). Together, these findings indicate that vestibular signals contribute fundamentally to embodied self-representation and agency, processes that are central to distinguishing self-generated from externally generated experiences. Yet their contribution to self-other processing in memory, particularly source and reality monitoring, remains largely unexplored.

Dissociative experiences provide a clinically relevant example of disrupted self-other processing at the embodied and cognitive levels. Depersonalisation and derealisation (DPDR) involve profound alterations in self-experience, including diminished agency, detachment from one’s body or mental states, and a reduced sense of ownership over experiences.

Depersonalisation can also alter the phenomenology of autobiographical memory, whereby individuals may feel detached from episodic memories, as though they had not been personally involved in those events (Sierra & David, 2011). Such reports suggest impairments in attributing experiences to the self, a process closely related to internal source monitoring and reality monitoring. Evidence indicates that vestibular dysfunction is associated with DPDR symptoms (Elyoseph et al., 2023), and in patients with vestibular dysfunction, caloric stimulation reproduced previously experienced symptoms during their vestibular impairment (Sang et al., 2006). These findings support the view that multisensory signals, including vestibular input, contribute to maintaining a coherent embodied self-model that supports accurate attribution of experiences across perception, action, and memory and have implications for the cognitive understanding of dissociative experiences.

To test whether vestibular signals contribute to the attribution of the origin of experiences, the current study used noisy galvanic vestibular stimulation (nGVS) to modulate vestibular input during an episodic memory task assessing source and reality monitoring. We also included both positive and threatening words to examine whether emotional content influenced source and reality monitoring. This design allowed us to test whether perturbing vestibular self-signals influences both the subjective experience of selfhood and the accuracy of attributing experiences to their source. Dissociative experiences were additionally assessed to examine whether alterations in embodied self-processing relate to memory attribution. We used both state and trait measures of DPDR, enabling us to test both the effects of nGVS on transient feelings of DPDR, as well as the impact of higher lifetime occurrence of DPDR on source and reality monitoring.

This design allowed us to test whether perturbing vestibular self-signals influences both the subjective experience of selfhood and the accuracy of attributing experiences to their source.

We hypothesised that active nGVS, relative to sham stimulation, would increase self-reported state DPDR symptoms and reduce both source- and reality-monitoring accuracy. We further predicted that greater trait DPDR history would be associated with poorer source- and reality-monitoring performance. Finally, we examined whether stimulus valence moderated the effects of nGVS on monitoring performance.

## 2. Methods

The study hypotheses, design, exclusion criteria, and analysis plan were preregistered on the Open Science Framework (https://osf.io/d29fe).

### 2.1 Participants

A total of 60 undergraduate students (58.33% female, *M*_age_ = 19.53, *SD_age_* = 1.75) from the University of Kent were recruited through the university’s research participation scheme in exchange for course credit. An a priori power analysis indicated that, with 30 participants per group (N = 60), the study would have approximately 80% power to detect a Group × Within-subject medium-large interaction effect of Cohen’s *f* = .37 at α = .05. Participants were pseudo-randomly assigned to either the active (real) or sham (placebo) stimulation conditions, with *N =* 30 in each group. The groups were matched for age and gender. Participants in the sham condition had a mean age of 19.87 years (SD = 2.01, range = 18–27), while participants in the active nGVS condition had a mean age of 19.20 years (SD = 1.40, range = 18–24). In the sham condition, 17 participants identified as female and 13 as male, while in the active nGVS condition, 18 participants identified as female and 12 as male. Inclusion criteria were: (a) aged 18-35 years, (b) normal or corrected-to-normal vision, and (c) fluency in English. Exclusion criteria included: (a) psychiatric or neurological diagnosis, (b) a history of epilepsy or family history of epilepsy, (c) use of psychiatric medication at the time of the experiment, (d) history of brain injury, concussion, seizures, or migraines, (e) presence of electrical medical equipment that could not be removed, (f) metal present in the head (excluding dental work), (g) skin sensitivities or conditions, and (h) prior participation in GVS studies. Participants who had previously participated in a GVS study were excluded as they would be aware from the electrode placement as to whether they were in the active or the sham condition. The study was conducted in accordance with the ethical principles of the Declaration of Helsinki and received ethical approval from the University of Kent Research Ethics Committee [Ethics ID: 202417056628678818]. All participants provided informed consent prior to participation.

### 2.2 Materials

#### 2.2.1 Demographic Measures

Participants were asked to report their age and gender.

#### 2.2.2 Risk Assessment and Adverse Effects Questionnaire

The risk assessment questionnaire was given to participants to ensure that it would be safe for them to participate in a study using GVS. The adverse effects measure was given following either real or sham stimulation to assess the extent of any side effects.

#### 2.2.3 Visual Analog Mood Scale (VAMS)

The Visual Analog Mood Scale (VAMS), developed by Folstein and Luria (1973), consists of 8 items designed to measure different mood states - sad, afraid, angry, tired, energetic, happy, confused, and tense. Each item is presented as a 100mm vertical line, with one end being indicative of an emotion and the other end as neutral, demonstrated with cartoon faces.

Participants are asked to place a mark along the line to indicate how they currently feel. We used a computerised version of the task, with a continuous scale composed of 100-points, these points were unlabelled to remain true to the original measure.

#### 2.2.4 Cambridge Depersonalisation Scale

Depersonalisation and derealisation history was assessed using the Cambridge Depersonalisation Scale (CDS; Sierra & Berrios, 2000). The CDS is a 29-item self-report measure that assesses the frequency and duration of depersonalisation/derealisation experiences over the past 6-months. In our study, we changed this to reflect lifetime occurrence, as clinically significant DPDR can occur transiently during times of high stress or as a result of drug-use (Hunter et al., 2003) and our goal was to understand whether our participants had ever had a subjective experience of DPDR and to what extremity/frequency this occurred. Items index alterations in bodily experience, perceptual anomalies, emotional numbing, and disturbances in the sense of agency and reality. For each item, respondents rate both how often the experience occurs (frequency) and how long it lasts (duration) using Likert-type scales; scores are summed to yield a total severity index. No items are reverse-coded. Higher scores indicate greater depersonalisation/derealisation severity. The Cambridge Depersonalisation Scale has demonstrated good reliability (Cronbach’s α = .89; split-half reliability = .92) and validity, correctly distinguishing patients with DSM-IV depersonalisation disorder from clinical comparison groups.

#### 2.2.5 DPDR State Inventory

To assess whether active nGVS resulted in any transient feelings of DPDR, state symptoms were assessed before and after the experiment using the Depersonalisation/Derealisation Inventory (DDI; Cox & Swinson, 2002). This questionnaire is designed to assess whether DPDR is occurring at the time of testing. The DDI is a 28-item self-report measure designed to quantify the frequency and intensity of anomalous experiences involving detachment from the self (depersonalisation) and unreality of the external world (derealisation). Items assess alterations in bodily experience, perceptual distortions, emotional numbing, and disruptions in the sense of agency or reality. Respondents rate each item on a Likert-type scale reflecting the extent to which they experience each phenomenon. No items require reverse-coding. Higher scores indicate greater severity of depersonalisation/derealisation symptoms. The Depersonalization-Derealization Inventory has demonstrated excellent internal consistency (α = .95), alongside evidence of validity through significant associations with anxiety, depression, somatic symptoms, panic severity, and feelings of unreality.

#### 2.2.6 Noisy Galvanic Vestibular Stimulation (nGVS)

A non-invasive single-channel direct current stimulator (DC-Stimulator Plus, NeuroConn) delivered nGVS. For active nGVS, bilateral noisy low-frequency stimulation was applied with the following parameters: an amplitude range of -1750 uA to +1750 uA, no offset, a fade-in time of 5 seconds, and a duration of 600 seconds. We selected these parameters as previous research has shown nGVS in this range to downweight vestibular signals in VR studies (Weech, Wall, & Barnett-Cowan, 2020). Carbon rubber electrodes (3 cm x 3 cm) with a layer of conductive gel were used to ensure effective stimulation. The same parameters were applied for sham stimulation, but the electrodes were positioned on the neck to produce similar skin sensations without affecting the vestibular system, see Figure 1 for positioning of electrodes across both conditions. The study was single-blinded, as the experimenter had to know which stimulation condition the participants were in to correctly apply the electrodes. To assess the success of blinding, participants were asked at the end of the session to guess whether they received active or sham stimulation.

**Figure 1.**
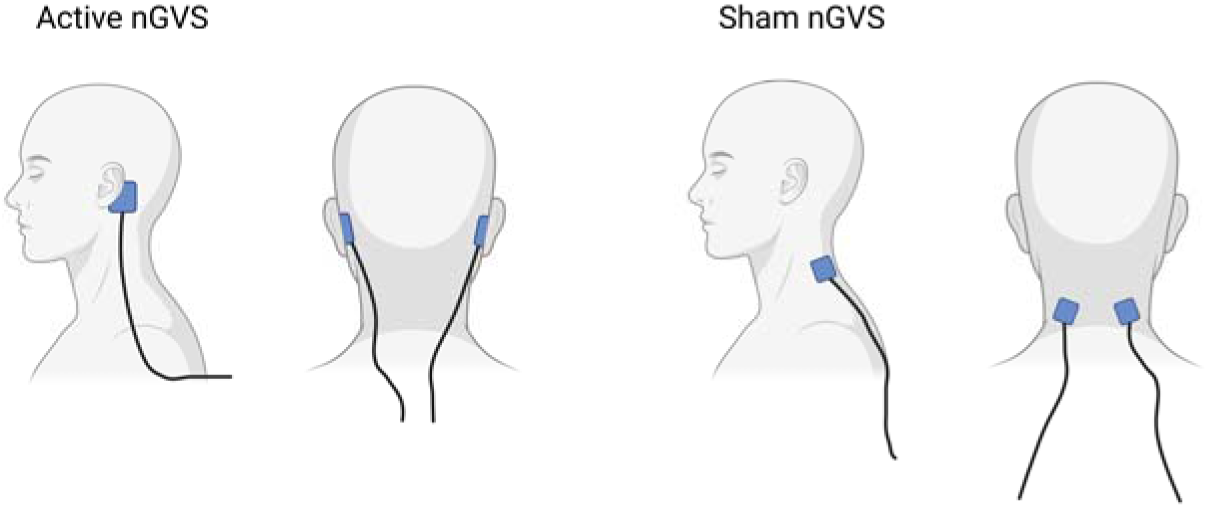
Positioning of nGVS electrodes in both the sham and the active conditions. *Note.* This schematic, created using BioRender, illustrates the electrode configuration used in the present study.

#### 2.2.7 Source-Reality Monitoring in Episodic Memory Task

Words were presented and encoded under four conditions: spoken aloud by the experimenter, spoken aloud by the participant, imagined as spoken by the participant, or imagined as spoken by the experimenter in a pseudorandomised order. The reality component indexed whether the event was externally generated (overt speech) or internally generated (imagined speech), whereas the source component indexed the agent to whom the word was attributed at encoding (self vs. experimenter), irrespective of reality status. Words were shown on the screen for 5-seconds; overall, the encoding task lasted for 6 minutes and 40 seconds. After encoding all 80 words, participants completed an unrelated 15-minute filler task (not reported here), followed by a memory test requiring cued-recall of both the encoding context (spoken vs. imagined) and the agent (self vs. experimenter). Stimuli were drawn from Warriner et al. (2013) and categorised as positive or negative according to valence ratings above or below 5, while being matched across categories on arousal. The experimenter was always female. Please see Figures 2 and 3 for representations of experimental trials and of the experimental set-up, respectively.

**Figure 2.**
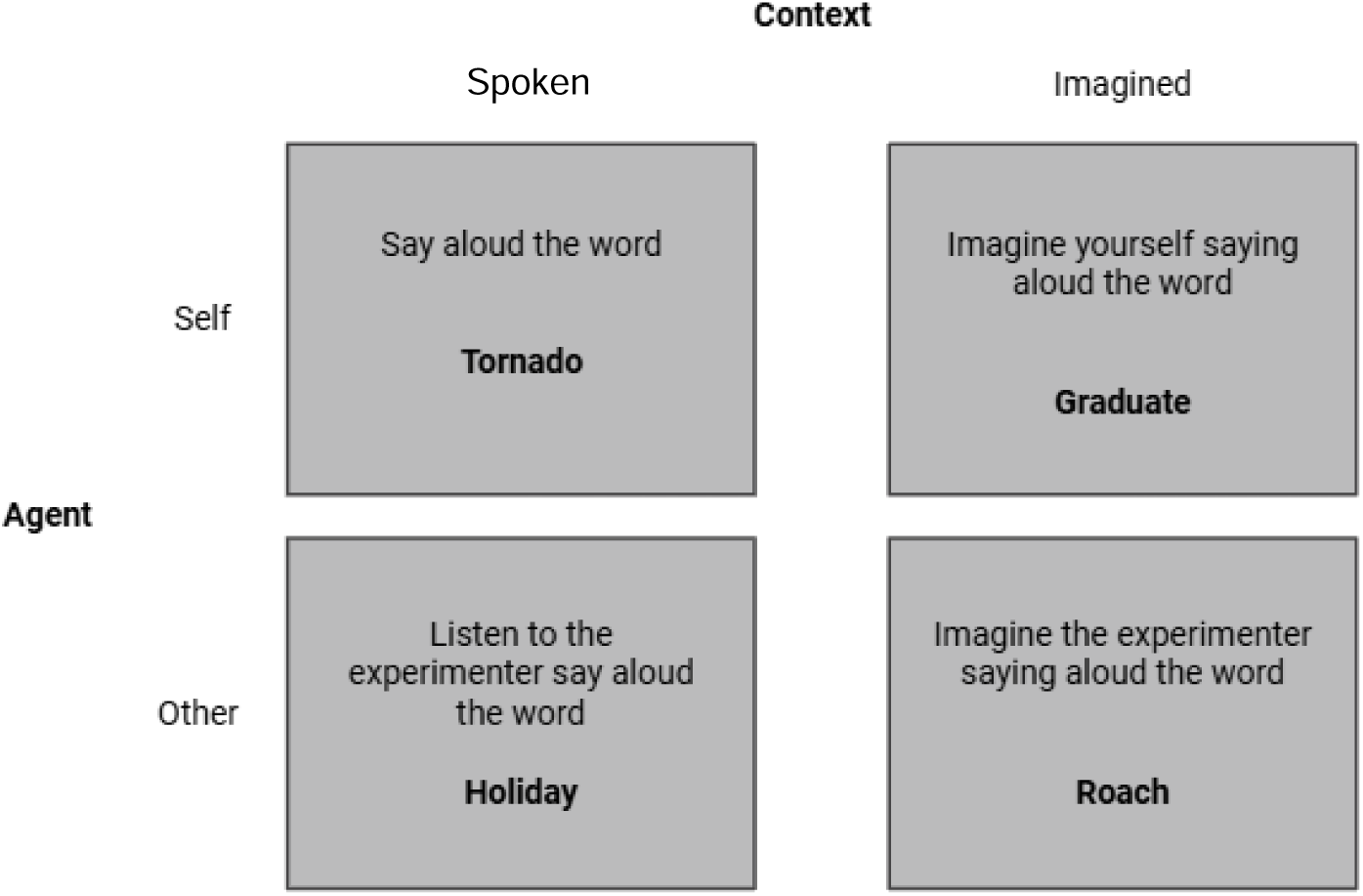
Visual representation of the experimental trials.

**Figure 3.**
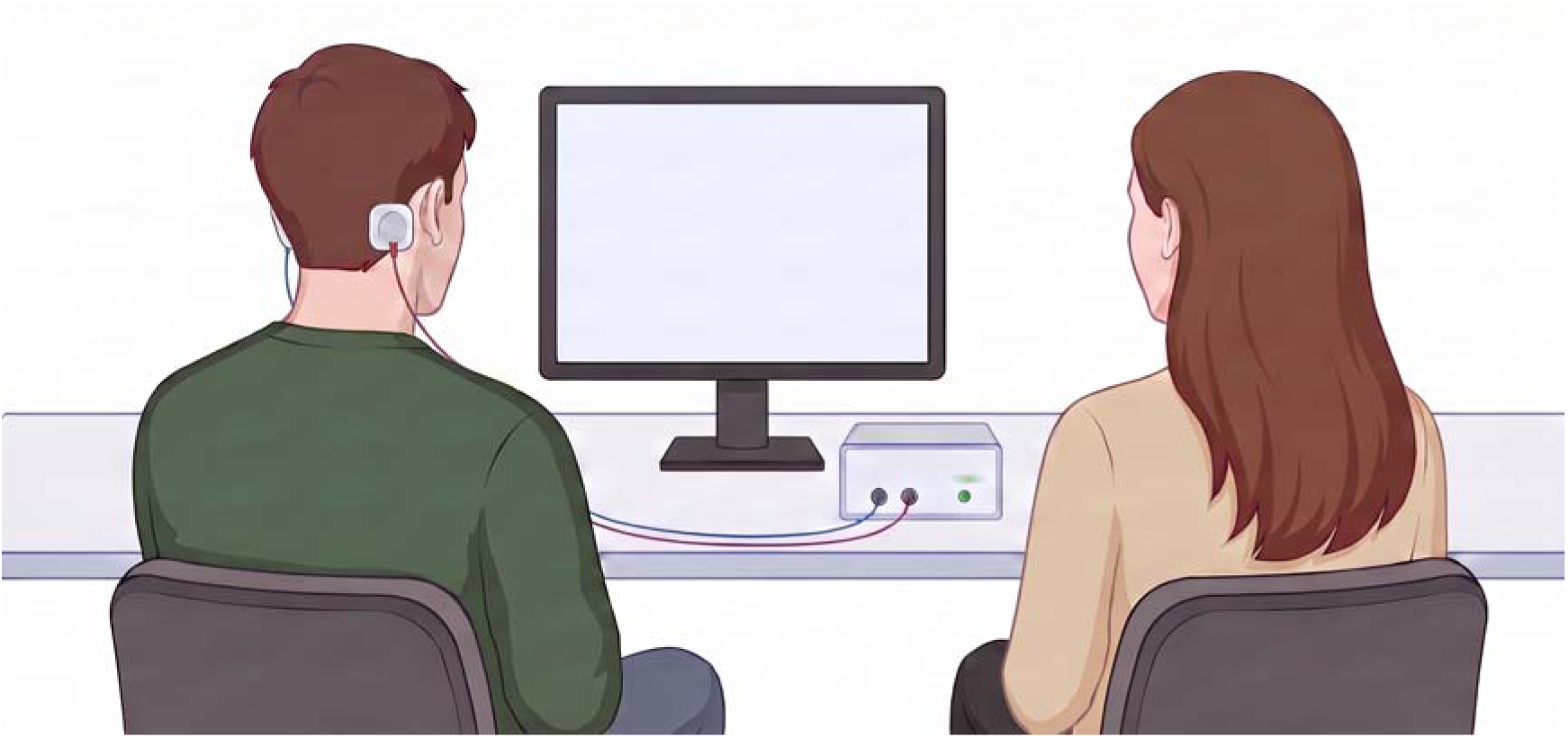
Visual representation of the experimental set-up. *Note.* This image is a mock-up of our actual experimental set-up, created in BioRender.

### 2.3 Design

The current study employs a mixed ANOVA design, with comparable analyses throughout each relevant DV (Reality and Source). For Reality, a 2x2x2 (Stimulation: sham, active x Agent: self, other x Valence: positive, threat) mixed repeated measures ANOVA was conducted on sensitivity measures (d-prime and criterion; see below for details), with Stimulation as a between-subjects factor and Agent and Valence as within-subject factors. For Source, a 2x2x2 (Stimulation: sham, active x Context: spoken, imagined x Valence: positive, negative) mixed repeated-measures ANOVA was conducted on sensitivity measures, with Stimulation as a between-subjects factor and Context and Valence as within-subjects factors.

#### 2.3.1 Signal-detection measures

The preregistered primary outcomes were discrimination sensitivity d-prime (d′) and response criterion (c). For each analysis, one source category was designated as the signal and the alternative source category formed the noise distribution.

For source monitoring, d′ quantified discrimination between self- and experimenter-generated items separately within each Context × Valence condition. Self-generated items were designated as the signal. Hits were therefore defined as self responses to self-generated items, whereas false alarms were defined as self responses to experimenter-generated items.

Experimenter responses to self-generated items constituted misses, and experimenter responses to experimenter-generated items constituted correct rejections.

For reality monitoring, d′ quantified discrimination between spoken and imagined items separately within each Agent × Valence condition. Spoken items were designated as the signal. Hits were therefore defined as spoken responses to spoken items, whereas false alarms were defined as spoken responses to imagined items. Imagined responses to spoken items constituted misses, and imagined responses to imagined items constituted correct rejections. See Table 1.

**Table 1.** Classification of responses used to calculate signal-detection measures.

|  | **True item category** | **Participant response** | **Classification** |
| --- | --- | --- | --- |
| **Source Monitoring** | Self | Self | Hit |
|  | Self | Experimenter | Miss |
|  | Experimenter | Self | False alarm |
|  | Experimenter | Experimenter | Correct rejection |
| **Reality Monitoring** | Spoken | Spoken | Hit |
|  | Spoken | Imagined | Miss |
|  | Imagined | Spoken | False alarm |
|  | Imagined | Imagined | Correct rejection |

Hits *(H)* and false alarm *(F)* rates were calculated as:

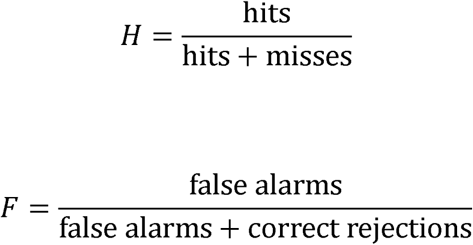

To avoid infinite values arising from hit or false-alarm rates of zero or one, the loglinear correction was applied to all condition-level rates:

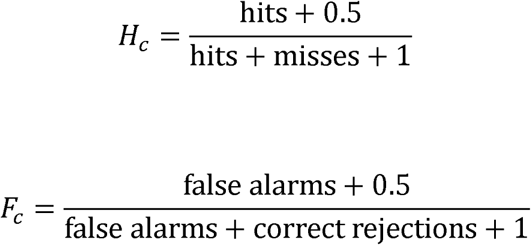

Discrimination sensitivity was then calculated as:

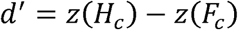

Higher values of d′ indicate greater sensitivity in discriminating the two source categories, whereas a value of zero indicates chance-level discrimination.

Response criterion was calculated as:

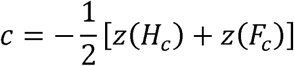

Criterion values indicate the tendency to favour one response independently of discrimination sensitivity. Positive values of c indicate conservative use of the designated signal response, whereas negative values indicate liberal use of that response. Thus, positive values reflected a conservative tendency to make a self-response in the source-monitoring analysis and a conservative tendency to make a spoken response in the reality-monitoring analysis.

#### 2.3.2 Trait DPDR and source and reality monitoring performance

Exploratory linear regression analyses were conducted to examine whether trait depersonalisation/derealisation (DPDR) history was associated with overall source- and reality-monitoring performance and whether it moderated the effects of vestibular stimulation. To maximise power for these exploratory analyses, sensitivity (d′) and response criterion (c) scores were averaged across the within-subject task conditions, yielding one overall d′ score and one overall criterion score for each participant for the source- and reality-monitoring tasks. Separate regression models were fitted for source-monitoring sensitivity, source-monitoring criterion, reality-monitoring sensitivity, and reality-monitoring criterion. Each model included stimulation condition (active nGVS vs. sham), mean-centred trait DPDR history, and the Stimulation × DPDR interaction as predictors. Stimulation condition was included to account for the primary experimental manipulation while evaluating the unique contribution of trait DPDR. As these analyses were exploratory and were not specified in the preregistration, findings were interpreted cautiously.

#### 2.3.3 Effects of nGVS on state DPDR

To examine whether nGVS influenced state depersonalisation/derealisation (DPDR) symptoms, a 2 × 2 mixed-design analysis of variance (ANOVA) was conducted with Time (pre-experiment, post-experiment) as a within-subjects factor and Stimulation (active nGVS, sham) as a between-subjects factor. Total state DPDR scores served as the dependent variable. The analysis tested the main effects of Time and Stimulation, as well as the Time × Stimulation interaction, to determine whether changes in state DPDR symptoms across the experimental session differed between stimulation conditions.

### 2.4 Procedure

Participants first completed a safety screening to confirm eligibility and absence of contraindications for GVS. Eligible participants received an information sheet and provided written informed consent. They then completed a demographic questionnaire (age, gender, ethnicity, hand dominance), the VAMS, the Cambridge Depersonalisation Scale, and the Depersonalisation/Derealisation Inventory. Electrodes were subsequently positioned according to condition: behind the ears for active stimulation targeting the vestibular system, or on the back of the neck for sham stimulation. Participants were informed about potential sensations and completed the cognitive task described above. Following the task, participants completed the VAMS and the DDI again, an adverse effects measure, and were asked to guess which stimulation condition they were in. Participants were then debriefed and reminded of their right to withdraw their data.

## 3. Results

The preregistration specified separate mixed ANOVAs examining source- and reality-monitoring performance (d’ and c), alongside analyses of state and trait DPDR (https://osf.io/d29fe). All post-hoc tests were Bonferroni-corrected separately within each family of related comparisons, and the corrected p-values are reported throughout.

Two participants performed with d’ < 0 during noisy GVS, suggesting extreme effects of stimulation or a failure to understand the task requirements. All analyses were re-calculated with these outliers removed with no significant changes to the outcomes. Therefore, for completeness, the followings analyses include the two outlying participants.

56.7% of participants (34/60) correctly identified the Stimulation condition (active vs. sham), but this did not differ significantly from chance (binomial test, two-tailed *p* = .36), indicating successful blinding. A separate binomial test indicated that participants were more likely to guess “real stimulation” than “sham stimulation,” with 40/60 responses indicating “real,” *p* = .013. This suggests a significant bias toward endorsing real stimulation above chance (50%).

A chi-square test indicated that gender distribution did not differ significantly across stimulation conditions, χ²(1, N = 60) = 0.07, *p* = .793. Additionally, there was no significant difference in age between stimulation groups, *t*(51.73) = 1.49, *p* = .142, *d* = 0.39.

Separate 2 (Time: pre, post) × 2 (Stimulation: sham, active nGVS) mixed ANOVAs were conducted for positive and negative mood. For positive mood, there was a significant main effect of Time, *F*(1, 58) = 5.21, *p* = .026, ηp² = .082, reflecting a reduction in mean positive mood from pre-task (*M* = 31.68, *SD* = 17.74) to post-task (*M* = 28.24, *SD* = 18.14), see Table 2. The main effect of Stimulation, *F*(1, 58) = 0.06, *p* = .803, and the Time × Stimulation interaction, *F*(1, 58) = 0.04, *p* = .847, were not significant.

**Table 2.** VAMS Pre and Post Scores and Adverse Effects.

| **Measure** | **Time** | **Sham nGVS**<br>***M (SD)*** | **Active nGVS,**<br>***M (SD)*** |
| --- | --- | --- | --- |
| **Positive Mood** | Pre | 32.38 (16.63) | 30.98 (19.05) |
|  | Post | 28.65 (19.15) | 27.83 (17.40) |
| **Negative Mood** | Pre | 19.12 (14.37) | 16.72 (10.04) |
|  | Post | 16.93 (13.83) | 12.38 (8.78) |
| **Adverse Effects** | Post | 3.80 (3.16) | 3.47 (2.92) |

*Note.* Positive mood scores were averaged across two positive-affect items, whereas negative mood scores were averaged across six negative-affect items. Adverse effects were assessed following stimulation.

For negative mood, there was also a significant main effect of Time, *F*(1, 58) = 14.43, *p* < .001, ηp² = .199, with mean negative mood decreasing from pre-task (*M* = 17.92, *SD* = 12.35) to post-task (*M* = 14.66, *SD* = 11.72), see Table 2. Neither the main effect of Stimulation, *F*(1, 58) = 1.37, *p* = .247, nor the Time × Stimulation interaction, *F*(1, 58) = 1.57, *p* = .216, was significant. Thus, although both positive and negative mood declined over the experimental session, there was no evidence that active nGVS differentially affected either dimension of mood.

There was no significant difference in reported adverse effects between stimulation conditions, *t*(57.66) = 0.43, *p* = .673, *d* = 0.11, see Table 2.

### 3.1 Source Monitoring

#### 3.1.1 Sensitivity (d’)

There was a significant main effect of Stimulation, *F*(1, 58) = 5.24, *p* = .026, ηG² = .044, such that source sensitivity was higher in the sham condition (*M* = 1.02, *SD* = 0.44) than in the active condition (*M* = 0.70, *SD* = 0.44). See Table 3 and Figure 4a.

**Figure 4.**
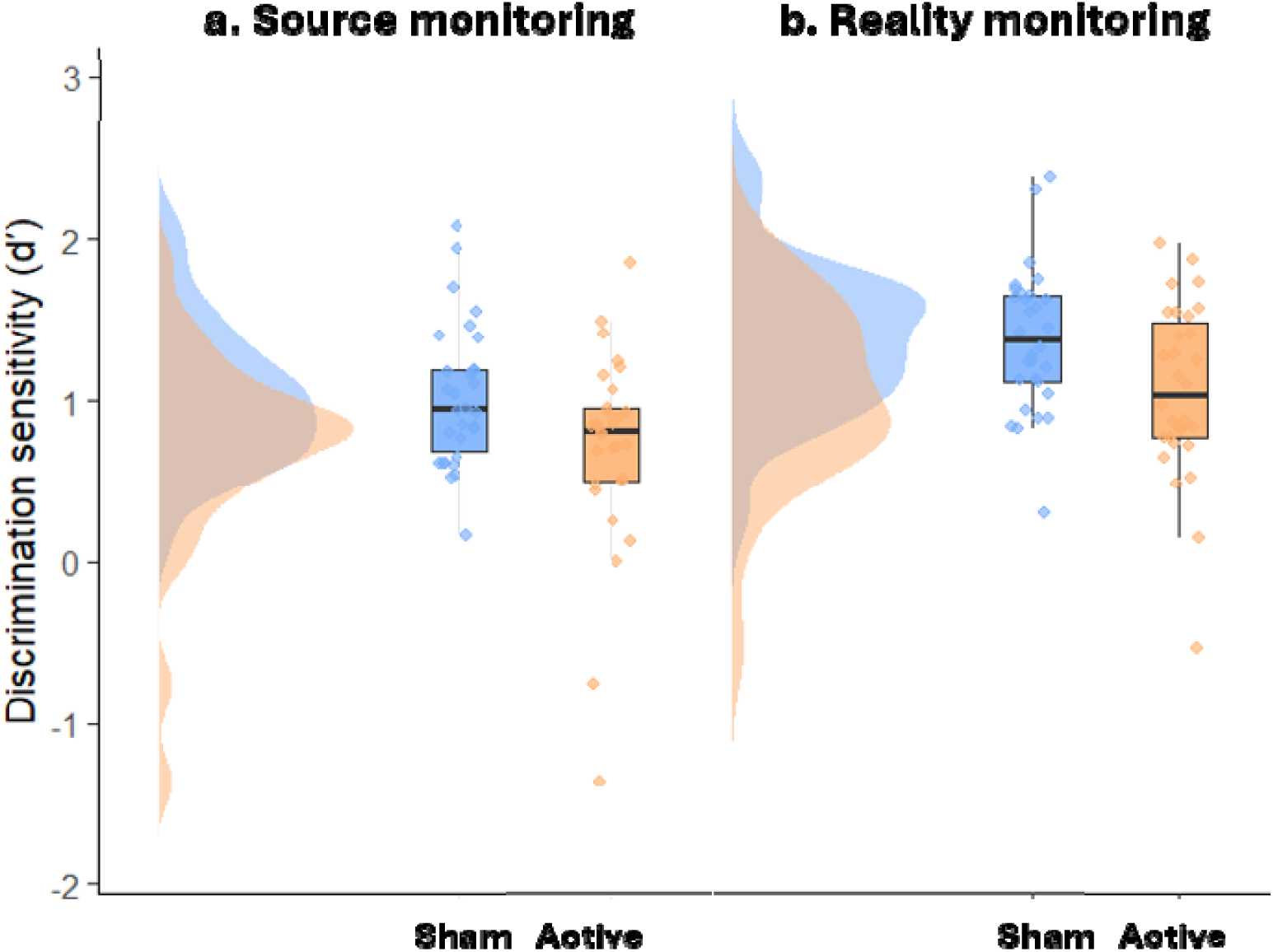
Main Effect of nGVS on Source and Reality Monitoring Sensitivity. *Note.* Points represent individual participants’ mean d’ scores, boxplots show the median and interquartile range, and shaded distributions show the density of scores within each stimulation condition. All observations are displayed. Analyses remain significant after removal of potential outliers (d’ < 0).

**Table 3.**
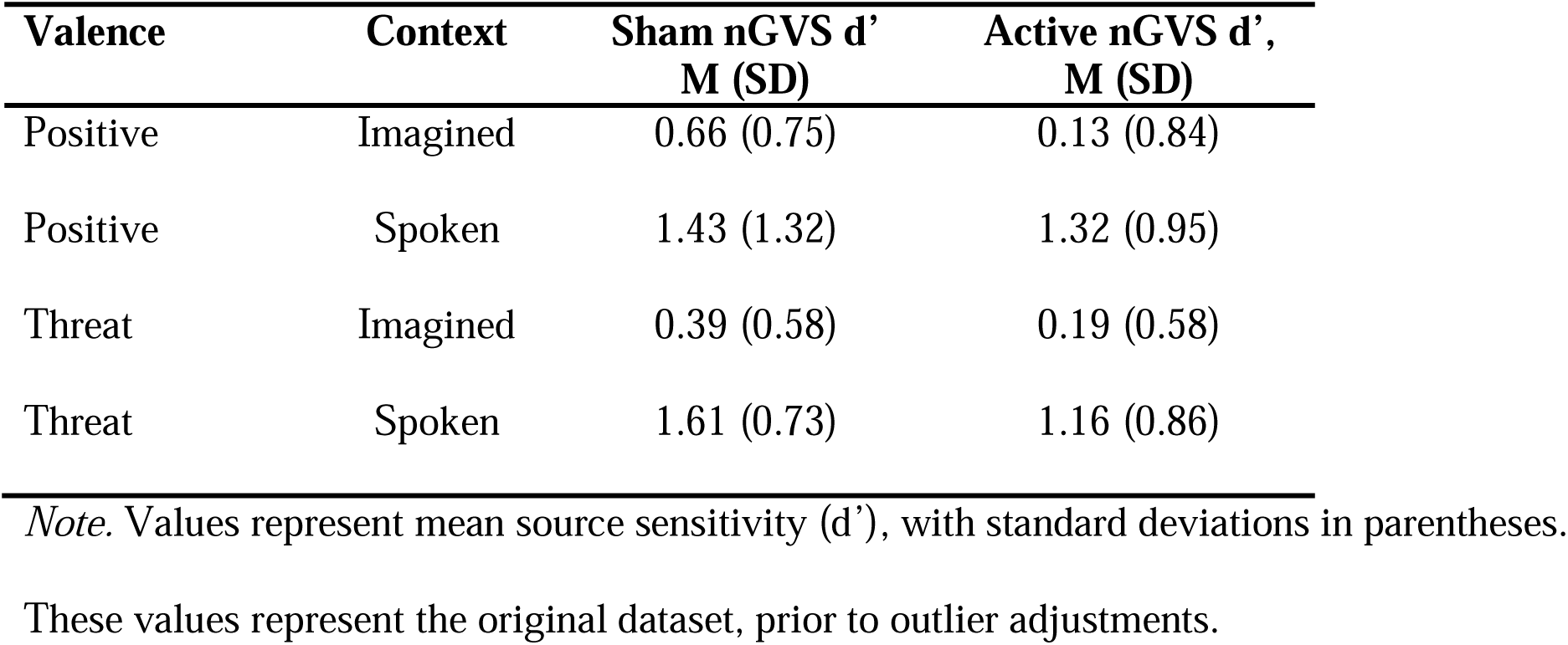
Descriptive Statistics for Source Monitoring Sensitivity Across all Conditions.

| **Valence** | **Context** | **Sham nGVS d'**<br>**M (SD)** | **Active nGVS d',**<br>**M (SD)** |
| --- | --- | --- | --- |
| Positive | Imagined | 0.66 (0.75) | 0.13 (0.84) |
| Positive | Spoken | 1.43 (1.32) | 1.32 (0.95) |
| Threat | Imagined | 0.39 (0.58) | 0.19 (0.58) |
| Threat | Spoken | 1.61 (0.73) | 1.16 (0.86) |

*Note.* Values represent mean source sensitivity ( $d'$ ), with standard deviations in parentheses.
These values represent the original dataset, prior to outlier adjustments.

A significant main effect of Context was also observed, *F*(1, 58) = 130.44, *p* < .001, ηG² = .325, with higher accuracy for spoken trials (*M* = 0.34, *SD* = 0.57) than for imagined trials (*M* = 0.34, *SD* = 0.57). The main effect of Valence was not significant, *F*(1, 58) = 0.34, *p* = .561, ηG² = .001.

In terms of interaction effects, the Stimulation × Context interaction was not significant, *F*(1, 58) = 0.21, *p* = .648, ηG² = .001, nor was the Stimulation × Valence interaction, *F*(1, 58) = 0.00, *p* = .944, ηG² < .001, or the Context × Valence interaction, *F*(1, 58) = 0.70, *p* = .405, ηG² = .001. However, the three-way Stimulation × Context × Valence interaction was significant, *F*(1, 58) = 6.28, *p* = .015, ηG² = .013. See below Figure 5.

**Figure 5.**
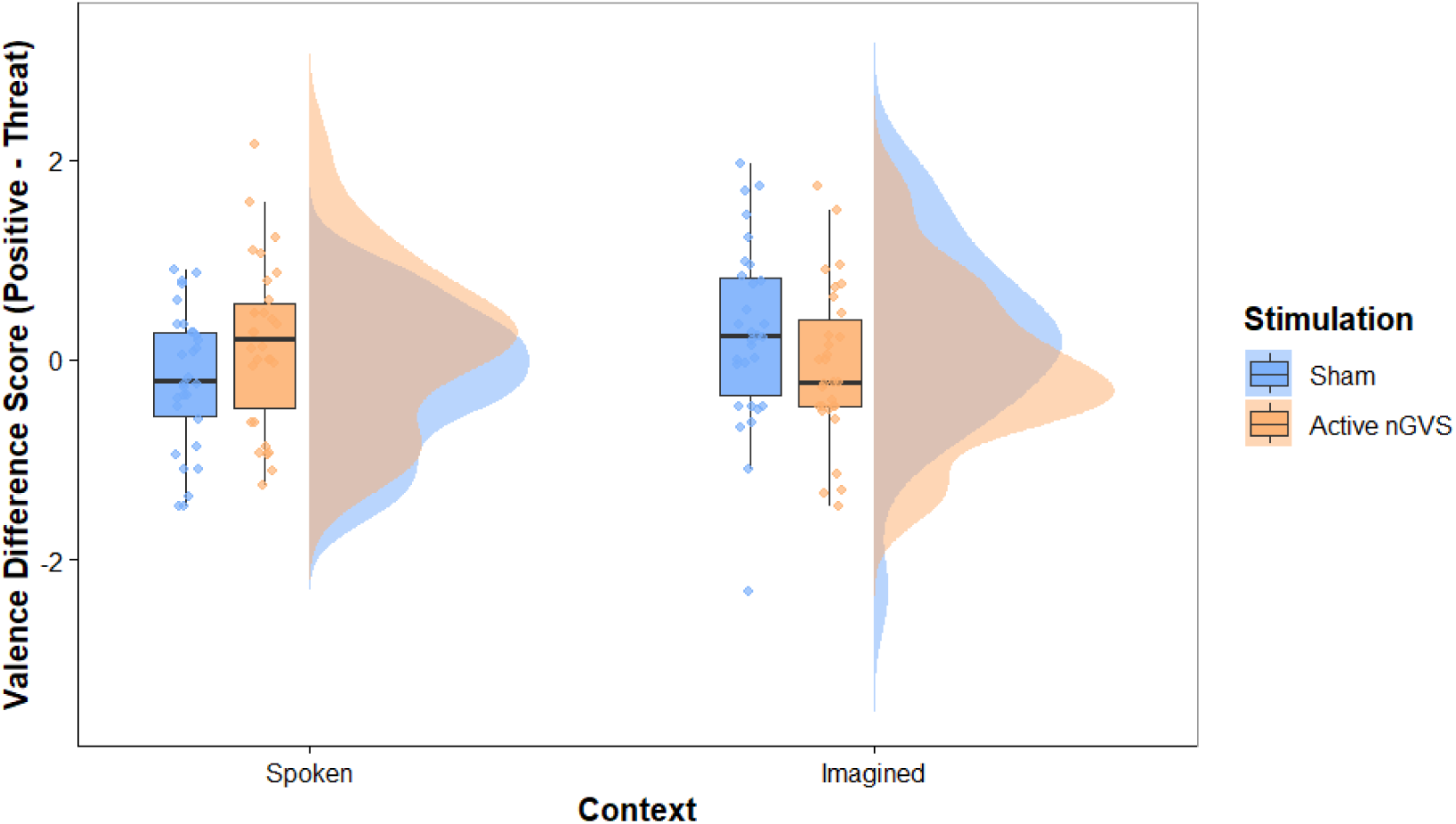
Interaction Effect between nGVS, Context, and Valence on Source Monitoring Sensitivity. *Note*. The difference score represents the valence effect on discrimination sensitivity (d′) and was calculated as d′ for positive words minus d′ for threat words. Positive scores indicate greater sensitivity for positive than threat words, whereas negative scores indicate greater sensitivity for threat than positive words.

To characterise the significant Stimulation × Context × Valence interaction, the Context × Valence interaction was examined separately within each stimulation condition. Under sham stimulation, the Context × Valence interaction was significant before correction, *F*(1, 29) = 4.68, *p* = .039, but did not survive Bonferroni correction, *p*_adj_ = .078. Under active nGVS, the interaction was not significant using either the uncorrected, *F*(1, 29) = 1.73, *p* = .199, or Bonferroni-adjusted criterion, *p*_adj_ = .398. The sham interaction was subsequently explored using simple-effects analyses to characterise the pattern underlying the omnibus interaction. However, because the stimulation-specific interaction did not survive correction for multiple comparisons, these simple effects should be interpreted cautiously. To further characterise the interaction during sham stimulation, paired-samples *t*-tests compared positive and threat words separately for imagined and spoken trials. There was no significant difference in source-monitoring sensitivity between positive and threat words for imagined trials, *t*(29) = −1.61, *p* = .119.

Similarly, no significant valence effect was observed for spoken trials, *t*(29) = 1.46, *p* = .155. In contrast, source-monitoring sensitivity was significantly lower for imagined than spoken trials for both threat words, *t*(29) = −6.87, *p* < .001, and positive words, *t*(29) = −4.80, *p* < .001.

Although the individual simple comparisons did not reach significance, the significant omnibus interaction indicates that the influence of valence differed across imagined and spoken contexts. Inspection of the means suggests that this differential pattern was attenuated during imagined and reversed during spoken trials following active nGVS.

#### 3.1.2 Response criterion (c)

There was no main effect of Stimulation on source criterion, F(1, 58) = 0.12, p = .734, ηG² = .001. The main effects of Context, F(1, 58) = 3.67, p = .060, ηG² = .015, and Valence, F(1, 58) = 1.06, p = .307, ηG² = .004, were also not significant. There was, however, a significant Stimulation × Context interaction, F(1, 58) = 5.51, p = .022, ηG² = .022. In the sham condition, criterion was more conservative for imagined trials (M = 0.28) than for spoken trials (M = 0.08), t(29) = 3.12, p = .008; this difference was not present in the active condition (imagined = 0.15, spoken = 0.17), t(29) = −0.30, p = 1.000. Between-group comparisons were not significant within either context (corrected ps ≥ .276). The Stimulation × Valence, F(1, 58) = 0.35, p = .556, ηG² = .001, Context × Valence, F(1, 58) = 0.21, p = .648, ηG² < .001, and Stimulation × Context × Valence interactions, F(1, 58) = 1.80, p = .185, ηG² = .003, were not significant. See Table 4.

**Table 4.** Descriptive Statistics for Source-Monitoring Criterion (c) Across All Conditions.

| **Valence** | **Context** | **Sham nGVS**<br>**M (SD)** | **Active nGVS**<br>**M (SD)** |
| --- | --- | --- | --- |
| Positive | Imagined | 0.29 (0.40) | 0.17 (0.37) |
| Positive | Spoken | 0.14 (0.29) | 0.16 (0.36) |
| Threat | Imagined | 0.27 (0.38) | 0.12 (0.52) |
| Threat | Spoken | 0.01 (0.33) | 0.17 (0.43) |

*Note.* Values represent mean source-monitoring criterion (c) scores, with standard deviations in parentheses. Positive values indicate a conservative criterion for responding “self” (i.e., a bias towards responding “experimenter”). Values are prior to outlier adjustments.

### 3.2 Reality Monitoring

#### 3.2.1 Sensitivity (d’)

The main effect of Stimulation was significant, *F*(1, 58) = 5.95, *p* = .018, ηG² = .044, indicating greater reality-monitoring sensitivity in the sham condition (*M* = 1.38, *SD* = 0.44) than in the active condition (*M* = 1.07, *SD* = 0.54). See Table 5 and Figure 4b.

**Table 5.** Descriptive Statistics for Reality Monitoring Sensitivity Across all Conditions.

| **Valence** | **Agent** | **Sham nGVS d'**<br>**M (SD)** | **Active nGVS d'**<br>**M (SD)** |
| --- | --- | --- | --- |
| Positive | Self | 1.55 (0.59) | 1.15 (0.86) |
| Positive | Experimenter | 1.13 (0.74) | 0.91 (0.66) |
| Threat | Self | 1.64 (0.80) | 1.37 (0.80) |
| Threat | Experimenter | 1.20 (0.68) | 0.82 (0.79) |

*Note.* Values represent mean source-monitoring sensitivity ( $d'$ ) scores, with standard deviations in parentheses. These values represent the original dataset, prior to outlier adjustments.

The main effect of Agent on d’ was also significant, *F*(1, 58) = 20.65, *p* < .001, ηG² = .073, showing greater sensitivity for self-referential trials (*M* = 1.43, *SD* = 0.66) compared to experimenter-referential trials (*M* = 1.02, *SD* = 0.58).

The main effect of Valence was not significant, *F*(1, 58) = 0.73, *p* = .397, ηG² = .002, indicating no significant difference between positive and threat trials overall.

No significant interactions were found between Stimulation and Agent, F(1, 58) = 0.03, p = .931, ηG² < .001; Stimulation and Valence, F(1, 58) = 0.01, p = .931, ηG² < .001; Agent and Valence, F(1, 58) = 1.25, p = .268, ηG² = .003; or the three-way interaction between Stimulation, Agent, and Valence, F(1, 58) = 0.85, p = .360, ηG² = .002.

#### ·***3.2.1*** Response Criterion (c)

There was a significant main effect of Agent on criterion, F(1, 58) = 13.69, p < .001, ηG² = .032. Criterion was more conservative for experimenter-referential trials (M = 0.43, SD = 0.35) than for self-referential trials (M = 0.29, SD = 0.32). The main effects of Stimulation, F(1, 58) = 1.47, p = .231, ηG² = .014, and Valence, F(1, 58) = 0.96, p = .331, ηG² = .002, were not significant. Neither were the Stimulation × Agent, F(1, 58) = 2.22, p = .142, ηG² = .005, Stimulation × Valence, F(1, 58) = 0.38, p = .538, ηG² = .001, Agent × Valence, F(1, 58) = 0.23, p = .630, ηG² = .001, or three-way interactions, F(1, 58) = 0.03, p = .865, ηG² < .001. See Table 6.

**Table 6.** Descriptive Statistics for Reality-Monitoring Criterion (c) Across All Conditions.

| Valence | Agent | Sham, M (SD) | Active, M (SD) |
| --- | --- | --- | --- |
| Positive | Self | 0.38 (0.42) | 0.26 (0.37) |
| Positive | Other | 0.45 (0.39) | 0.43 (0.29) |
| Threat | Self | 0.35 (0.40) | 0.17 (0.36) |
| Threat | Other | 0.45 (0.43) | 0.39 (0.45) |

*Note.* Values represent mean reality-monitoring criterion (c) scores, with standard deviations in parentheses. Positive values indicate a bias towards responding “imagined”. Values are prior to outlier adjustments.

### 3.3 Trait DPDR and source and reality monitoring performance

To examine whether trait depersonalisation/derealisation (DPDR) was associated with overall monitoring performance, exploratory linear regression models were conducted using averaged sensitivity (d′) and response criterion (c) scores across task conditions. Stimulation condition was included as a covariate in all models, together with centred DPDR history and the DPDR × Stimulation interaction.

For source monitoring, trait DPDR did not significantly predict either sensitivity (β = 0.006, *p* = .177) or response criterion (β = 0.003, *p* = .088). The interaction between DPDR and stimulation was also not significant for either sensitivity (β = −0.004, *p* = .463) or response criterion (β = −0.005, *p* = .070).

For reality monitoring, trait DPDR did not predict sensitivity (β = −0.004, *p* = .353), and there was no evidence that DPDR moderated the effect of stimulation (β = 0.005, *p* = .367).

However, higher trait DPDR significantly predicted a lower reality-monitoring response criterion (β = −0.006, *p* = .009), indicating a more liberal bias when judging stimuli as real. The DPDR × Stimulation interaction for response criterion was not significant (β = 0.006, *p* = .077).

### 3.4 State DPDR effects of nGVS

A 2 (Time: pre, post) × 2 (Stimulation: active nGVS, sham) mixed ANOVA was conducted to examine whether state DPDR symptoms changed over the course of the experiment and whether this change differed by stimulation condition. There was a significant main effect of Time, F(1, 58) = 4.64, p = .035, indicating that DPDR scores were higher post-experiment (M = 40.25, SD = 11.89) than at baseline (M = 37.85, SD = 11.99). However, the Time × Stimulation interaction was not significant, F(1, 58) = 0.56, p = .457, indicating that the increase in DPDR symptoms did not differ between the active nGVS and sham conditions. The main effect of Stimulation was also not significant, F(1, 58) = 0.42, p = .521. Please see Table 7 for the means and standard deviations. Together, these findings suggest that participants reported a small increase in state DPDR symptoms across the experimental session, but this increase was not specific to active nGVS.

**Table 7.** Change in DPDR Symptoms Across The Experimental Session.

| **Stimulation Group** | **Pre, *M* (*SD*)** | **Post, *M* (*SD*)** |
| --- | --- | --- |
| **Sham** | 37.33 (12.70) | 38.90 (12.67) |
| **Active nGVS** | 38.37 (11.42) | 41.60 (11.11) |

## 4. Discussion

In the present study, we assessed how perturbation of vestibular signals using noisy galvanic vestibular stimulation (nGVS) influenced source and reality attributions in episodic memory, as well as the moderating role of emotional valence. We also examined whether disruption of vestibular signalling induced symptoms of depersonalisation and derealisation (DPDR), and whether trait DPDR was associated with monitoring performance. Active nGVS significantly reduced sensitivity (d′) in both source and reality monitoring, providing causal evidence for the involvement of the vestibular system in contextual aspects of episodic memory. For source monitoring, this effect was qualified by an interaction between stimulation, context, and valence, with reductions in sensitivity particularly evident for imagined and threat-related information. Stimulation did not produce an overall shift in response criterion (c) in either task, although it altered the context-dependent criterion pattern in source monitoring. Contrary to our prediction that trait DPDR would be associated with poorer monitoring sensitivity, higher trait DPDR was instead associated with a more liberal reality-monitoring response criterion, while remaining unrelated to discrimination sensitivity. Finally, although dissociative experiences increased across the experimental session, this increase did not differ between active and sham vestibular stimulation.

The present findings support embodied accounts of episodic memory, which propose that remembering the origins of experiences depends not only on mnemonic representations but also on multisensory signals that contribute to bodily self-representation (Ianì, 2019; Tsakiris, 2017; Blanke, 2012). Source and reality monitoring require individuals to distinguish internally generated experiences from those originating in the external world, a process likely supported by the integration of vestibular, proprioceptive and visual signals that continuously update the sense of self. By perturbing one component of this system, nGVS may have reduced the reliability of bodily cues that normally help distinguish self-generated from externally generated experiences, making memory attribution less precise.

One plausible neural substrate for these computations is the temporoparietal junction (TPJ), a multisensory hub involved in integrating vestibular, visual, and proprioceptive information to support self-location, sense of agency (Pryke et al., 2025), and self-other distinction (Blanke, 2012; Decety & Lamm, 2007; Quesque & Brass, 2019). Beyond its role in bodily self-representation, the TPJ also contributes to episodic memory retrieval, especially the retrieval of contextual information associated with remembered events (Vilberg & Rugg, 2008; Rugg & Vilberg, 2013). Disrupting vestibular input may therefore reduce the precision of multisensory representations that are later used to attribute memories to their source. Although previous studies have largely examined suprathreshold galvanic vestibular stimulation, recent neuroimaging evidence indicates that nGVS also modulates activity within the temporoparietal network, including the supramarginal gyrus (Valdés et al., 2021). Rather than simply increasing or suppressing cortical activity, nGVS may therefore introduce noise into multisensory computations supporting memory attribution. This interpretation is consistent with the present findings, in which vestibular perturbation reduced the precision of both source and reality monitoring.

An alternative explanation is that vestibular perturbation produced a non-specific cognitive deficit. However, this interpretation is difficult to reconcile with the present pattern of results. Rather than uniformly reducing performance, nGVS selectively impaired source and reality monitoring while interacting with task context and emotional valence, suggesting a disruption of processes supporting memory attribution rather than a general reduction in cognitive function. This interpretation is broadly consistent with previous work demonstrating that vestibular stimulation can produce selective effects on cognition. For example, suprathreshold galvanic vestibular stimulation has been shown to impair perspective-taking while leaving other cognitive functions, including reaction time and mental rotation, relatively unaffected (Dilda et al., 2011). Although these studies employed different stimulation parameters from the present investigation, they nevertheless suggest that vestibular stimulation does not produce a uniform cognitive deficit. Future studies incorporating a broader cognitive test battery and directly comparing stimulation protocols will be important for establishing the specificity of these effects.

A novel and interesting finding was the selective impairment of source monitoring for threatening information following active nGVS. This finding extends evidence that vestibular and emotional processing interact bidirectionally. Preuss et al. (2015) found that vestibular motion discrimination improved when participants viewed negatively valenced, threat-relevant images relative to neutral and positive images, suggesting that emotional salience can sharpen vestibular processing when accurate bodily orientation may be particularly adaptive. Whereas the current study demonstrates that emotional information can modulate vestibular perception, the present findings suggest the opposite direction of influence, with perturbing vestibular input selectively disrupting memory for threatening information. This bidirectional relationship may reflect overlap between vestibular and fear-processing networks, including the anterior insula, anterior cingulate cortex, thalamus and temporoparietal regions (Neumann et al., 2023; Lotze et al., 2025). Importantly, this effect was observed only for imagined items. One possibility is that imagined threat relies more heavily on internally generated multisensory representations than spoken threat, because there are fewer external perceptual cues available at encoding. Under this account, perturbing vestibular input may disproportionately reduce the precision of internally constructed representations supporting source attribution for imagined threat. Nevertheless, this mechanistic interpretation remains tentative and should be examined directly in future neuroimaging research.

Across stimulation conditions, reality-monitoring sensitivity was higher for self- than experimenter-generated speech, while source-monitoring sensitivity was higher for spoken than imagined speech. The latter finding is consistent with Kwon et al. (2022), who similarly reported poorer attribution for imagined than perceived events, suggesting that internally generated experiences provide fewer distinctive cues for subsequent source judgements. In contrast, the greater sensitivity for self-generated speech differs from Kwon et al.’s finding of poorer attribution for self-performed actions. This discrepancy may reflect differences in the information available to support source judgements. Self-generated speech provides multiple cues to agency, including speech production, auditory feedback and speaker identity, whereas Kwon et al.’s task involved visually similar object movements that offered fewer distinctive cues for distinguishing self- from other-generated actions. Together, these findings suggest that the influence of self-generation on source monitoring depends on the nature of the cues available at encoding.

The criterion analyses provide additional insight into the mechanisms underlying these effects by distinguishing mnemonic sensitivity from response tendencies. Consistent with the Source Monitoring Framework (Johnson & Raye, 1981; Johnson et al., 1993), source judgements depend not only on the information contained within a memory but also on the decision criteria used to evaluate that information. In the reality-monitoring task, participants adopted a more conservative criterion for judging experimenter-generated than self-generated speech as real, although the corresponding effect on discrimination sensitivity indicates that this difference cannot be explained by response bias alone. In source monitoring, nGVS did not produce an overall shift in criterion but eliminated the more conservative response criterion for imagined relative to spoken self-attributions observed in the sham group, suggesting that vestibular perturbation altered the context-dependent evaluation of memories without producing a general response bias.

Exploratory analyses showed that higher trait depersonalisation/derealisation (DPDR) was associated with a more liberal reality-monitoring response criterion, while remaining unrelated to discrimination sensitivity. Individuals reporting greater lifetime DPDR symptoms therefore appeared to adopt a lower threshold for judging experiences as having occurred in reality, despite preserved discrimination between imagined and spoken events. This suggests that vestibular perturbation and dissociative traits may influence memory attribution through different mechanisms: whereas nGVS primarily reduced mnemonic discrimination (d′), higher trait DPDR was associated with altered decision criteria during memory evaluation. Although this finding requires replication, it is consistent with the Source Monitoring Framework, which proposes that source-monitoring errors may arise from changes in either the quality of memory evidence or the criteria used to evaluate that evidence.

Unlike Sang et al. (2006), we found no evidence that vestibular perturbation increased self-reported depersonalisation/derealisation (DPDR). One likely explanation is that Sang et al. studied patients with vestibular disorders who already experienced chronic DPDR symptoms, whereas our sample comprised healthy participants receiving transient, largely imperceptible vestibular modulation. This interpretation is consistent with evidence that DPDR commonly accompanies chronic vestibular dysfunction (Elyoseph et al., 2023), suggesting that persistent or clinically significant disruption of vestibular processing may be necessary before subjective disturbances of self-experience emerge. More broadly, DPDR is likely to reflect the downstream consequences of prolonged disruption to multisensory self-processing rather than isolated perturbation of vestibular signals. Under this account, the cognitive changes observed here may represent an early stage of altered self-processing that is insufficient, on its own, to produce conscious dissociative symptoms.

Methodological differences between studies may also have contributed to the discrepant findings. Unlike Sang et al. (2006), the present study included a sham condition, revealing small increases in DPDR symptoms following both active and sham stimulation. This suggests that participation in an unusual experimental procedure or heightened attention to internal states may be sufficient to produce mild dissociative experiences (Schweden et al., 2018). Furthermore, CVS and nGVS differ fundamentally in their physiological effects. Whereas CVS produces strong, consciously perceived vestibular illusions, nGVS provides diffuse, largely imperceptible modulation of vestibular afferent activity. Because DPDR is inherently phenomenological, requiring conscious alterations in bodily or perceptual experience, the absence of overt vestibular sensations during nGVS may explain why vestibular perturbation impaired memory attribution without inducing subjective dissociative symptoms.

An important avenue for future research is to determine whether the conscious perceptual consequences of vestibular stimulation, rather than vestibular modulation itself, contribute to dissociative symptoms. Vestibular disorders such as vertigo and Meniere’s disease, which are characterised by erroneous self-motion perception and dizziness, have been associated with DPDR (Jáuregui-Renaud, 2015; Cento et al., 2026). One possibility is that sensory conflict between vestibular and other sensory cues, rather than vestibular noise alone, underlies these symptoms. Future studies could therefore employ stimulation techniques that evoke illusory self-motion, such as sinusoidal GVS, and directly relate subjective vestibular symptoms (for example dizziness or vection) to changes in DPDR.

A limitation of the present study is the absence of a direct physiological measure of vestibular modulation, such as postural sway or vestibulo-ocular reflex recordings. Consequently, although nGVS was administered using established protocols, it cannot be confirmed that vestibular processing was modulated to the same extent in every participant. In addition, the sample comprised young, healthy adults and was powered to detect medium-to-large effects, limiting both the generalisability of the findings and the ability to detect smaller effects, particularly for the higher-order interactions. Moreover, the exploratory analyses examining whether vestibular perturbation had greater effects on source and reality monitoring in individuals with higher levels of trait DPDR were underpowered. Although these analyses should therefore be interpreted cautiously, the observed trends warrant replication in larger samples. Future studies incorporating objective vestibular measures, larger samples and clinical populations will be important for establishing the reliability and generalisability of vestibular contributions to source and reality monitoring.

The present study provides novel evidence that vestibular self-signals contribute to the attribution of experiences in episodic memory. Active noisy galvanic vestibular stimulation impaired both source- and reality-monitoring sensitivity, supporting the view that accurate memory attribution depends, in part, on intact vestibular contributions to embodied self-representation. These impairments were particularly evident for imagined and threat-related information during source monitoring, suggesting that vestibular signals may be especially important when memory judgements rely on internally generated or emotionally salient representations. Although nGVS did not increase state depersonalisation/derealisation (DPDR) symptoms or interact with trait DPDR, exploratory analyses indicated that higher trait DPDR was associated with a more liberal reality-monitoring response criterion, suggesting that dissociative traits may influence memory attribution through changes in decision bias rather than mnemonic sensitivity. Together, these findings extend embodied accounts of episodic memory by demonstrating that vestibular perturbation disrupts memory discrimination, whereas dissociative traits appear to influence how memories are evaluated at retrieval. Future research should determine whether these effects generalise to clinical populations with chronic vestibular dysfunction or persistent DPDR and further investigate the neural mechanisms linking vestibular self-signals, bodily self-representation and episodic memory.

## Acknowledgements

The authors acknowledge all participants for their time and effort

## Data availability

All data will be available on the Open Science Framework (https://osf.io/d29fe).

